# Tissue resident colonic macrophages persist through acute inflammation and adapt to aid tissue repair

**DOI:** 10.1101/2025.09.19.677333

**Authors:** Lizi M. Hegarty, Gareth-Rhys Jones, Claire E. Adams, Rebecca M. Gentek, Gwo-Tzer Ho, Elaine Emmerson, Calum C. Bain

**Affiliations:** Centre for Inflammation Research, Institute for Regeneration and Repair, University of Edinburgh, 4-5 Little France Drive, Edinburgh, EH16 4UU, UK; Centre for Regenerative Medicine, Institute for Regeneration and Repair, University of Edinburgh, 4-5 Little France Drive, Edinburgh, EH16 4UU, UK; Centre for Immunobiology, School of Infection and Immunity, College of Medicine, Veterinary Medicine and Life Sciences, University of Glasgow, Glasgow, G12 9TA, UK; Centre for Reproductive Health, Institute for Regeneration and Repair, University of Edinburgh, 4-5 Little France Drive, Edinburgh, EH16 4UU, UK

**Keywords:** macrophages, colitis, inflammation, ontogeny, repair

## Abstract

Macrophages are crucial for the maintenance of intestinal homeostasis, are considered key proinflammatory effector cells during intestinal inflammation and are implicated in tissue repair following injury or inflammation. Whether these roles are attributed to distinct subsets of macrophages or if macrophages retain a degree of plasticity in the intestine remains poorly understood. Here, through a combination of single cell RNA sequencing, lineage-tracing and immunofluorescence imaging, we define three major subpopulations of murine, colonic macrophages on the basis of CD11c and CD163 expression. These macrophages occupy discrete anatomical niches and display distinct replenishment kinetics. They all accumulate during acute colitis and *Cx3cr1-*based fate mapping shows that they persist through to inflammation resolution. Moreover, marked transcriptional differences exist between the macrophages present in health and their counterparts in the post-inflammation environment, demonstrating that inflammation leads to transcriptional rewiring of the resident macrophages in a subset-specific manner. Intriguingly, there were minimal transcriptional changes between long-lived macrophages and their recently differentiated counterparts, indicating the environment exerted a greater influence than ontogeny or the time of residency on their functional states in resolution.

## Introduction

Macrophages play multiple roles to maintain homeostasis along the length and throughout the layers of the intestine[1], [2], [3]. This includes general housekeeping functions, such as the clearance of apoptotic and effete cells, but also more specialised functions, such as supporting epithelial renewal, blood vessel integrity, T cell maintenance, and regulation of the microbiome[1], [2], [3]. It is becoming increasingly clear that these functions are carried out by specific macrophage subsets which adopt phenotypic and functional specialisations in response to local environmental signals from their tissue microenvironment, and that bidirectional crosstalk of macrophages with other cells in their niche is essential for homeostasis[4], [5], [6]. In turn, the length of time spent in a particular tissue niche (‘time of residency’) is thought to impact macrophage differentiation/specialisation and restrict plasticity[7]. Thus, to fully understand macrophage function(s) in the intestine there is a great need to characterise their heterogeneity, location in the tissue and ‘time of residency’. Indeed, many previous studies, including ours, have focussed on understanding these parameters using mouse models[4], [8], [9], [10], [11], [12], [13], [14], [15]. We originally showed that CCR2-dependent, classical Ly6C^hi^ monocytes continually enter the colonic lamina propria to replenish mature macrophages[8], [9], [10]. Since then, there have been a number of studies showing intestinal macrophages in mice to be heterogeneous and describing the presence of long-lived macrophages in the intestine[4], [12], [16], [17], [18], [19]. However, there is a lack of consensus on how to define macrophage heterogeneity, with some studies defining subpopulations on the basis of Tim4 and CD4 expression[12], [20], and others using a combination of CCR2, CX3CR1, CD206, CD169 and/or CD121b[10], [12], [18], [19], [20].

As well as being considered ‘guardians of homeostasis’[21], monocytes and their macrophage progeny are implicated in inflammatory disorders of the intestinal tract, including inflammatory bowel disease (IBD), comprising Crohn’s disease and ulcerative colitis[1]. These chronic, debilitating conditions are characterised by periods of inflammation of unknown aetiology. While IBD pathogenesis is multifactorial, dysregulated monocyte/macrophage behaviour is a characteristic feature, with the accumulation of highly proinflammatory monocytic cells and their immediate progeny[9], [22], [23], [24], [25], [26]. Genetic or pharmacological depletion of monocytic cells in mice ameliorates experimental colitis, suggesting these cells are pathogenic[9], [11], [27]. In contrast, mature macrophages are considered to be terminally differentiated, refractory to proinflammatory signals and unable to respond in a similar manner during acute inflammation[11], [22], [25], [28]. However, if these mature, longer-lived macrophages persist through periods of inflammation and whether they adapt to aid inflammation resolution and tissue repair remains poorly understood. Understanding these processes is important given the relapsing, remitting nature of IBD, with 11.5% of all patients experiencing a clinically significant flare in their disease per year[29]. Moreover, many develop fibrotic disease in which excessive repair processes, including dysregulated monocyte/macrophage activity, is implicated[1], [30].

Here, we have used scRNAseq, lineage tracing, multiparameter immunofluorescence and flow cytometry to characterise the intestinal macrophage compartment in health and inflammation resolution, in mice. We show that in the mouse, expression of CD11c and CD163 allows partitioning of macrophages into transcriptionally and anatomically defined subsets with distinct replenishment kinetics in health. We show that these kinetics are affected in a subset-specific manner during and following inflammation. Moreover, we show that long-lived macrophages persist through periods of inflammation and adapt during resolution to aid repair, with distinct subsets playing discrete roles.

## Results & Discussion

### Expression of CD11c & CD163 defines colonic macrophage subsets

To characterise the colonic macrophage compartment and allow comparison between health and inflammation resolution in mice, we first performed single cell RNA-seq (scRNAseq) of live CD45^+^ CD11b/CD11c^+^ non-granulocytes from steady state colon using the 10X platform (**Supplementary** Fig. 1A-E). Cells of the monocyte/macrophage lineage were identified on the basis of their expression of *Csf1r*, *Adgre1* and/or *Fcgr1*, with other myeloid cells (e.g., conventional dendritic cells (cDC)) and contaminating cell types removed from subsequent analysis (**Supplementary** Fig. 1B, C). This revealed three clusters, which we defined as monocytes or mature macrophages on the basis of *Ly6c2* and *C1qa,* respectively (**Fig. 1A, B**). To increase the granularity of the mature, tissue-resident macrophage populations, we subclustered on the basis of *C1qa* and lack of *Ly6c2* to remove the monocytes. This led to 3 macrophage clusters being identified (**Fig. 1C**). Cluster 0 was defined by the expression of *Itgax*, *Dnase1l3*, *Il1r2*, *Mmp13,* and *Acp5* (Cluster 0), whereas Cluster 1 was defined by *Cd163, Folr2, Lyve1* and *Colec12 (***Fig. 1C, D**). The final cluster (Cluster 2) lacked expression of both *Itgax* and *Cd163* and was defined by its expression of *S100a6, Vcam1* and *Vim* (**Fig. 1C, D**). Interestingly, while *Mrc1* (encoding CD206) expression was a feature of Cluster 1 and has been shown to be coincident with *Cd163* in other tissues[31], expression was also evident in Clusters 0 and 2, albeit at lower levels (**Fig. 1D**).

**Figure 1.**
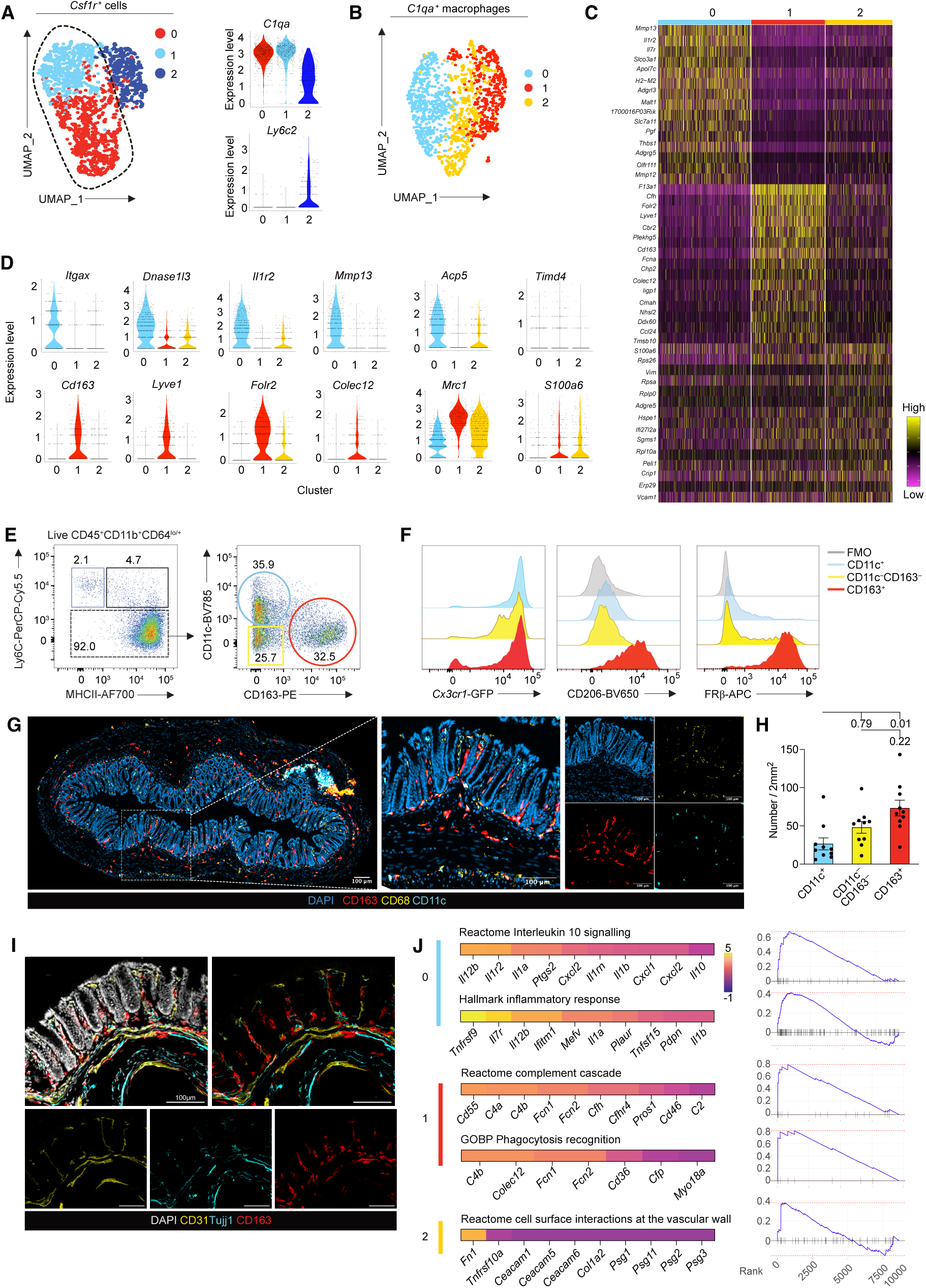
Expression of CD11c and CD163 defines distinct colonic macrophage subsets. **A.** UMAP projection of colonic *Csf1r*^+^ cells from naïve adult mice (as subclustered in Supplementary Fig. 1) and violin plots of selected cluster-defining genes. Cells from three mice were pooled for analysis. **B.** UMAP of sub-clustered (*C1qa*^+^*Ly6c2^−^*) macrophages. **C.** Heatmap displaying the top 15 cluster defining genes for each cluster of macrophages in B. **D.** Gene expression levels of selected cluster defining genes across the macrophage subsets. **E.** Representative flow cytometry plot of Ly6C vs MHCII expression by CD11b^+^CD64^lo/+^ cells to identify newly extravasated monocytes (Ly6C^+^MHCII^-^), intermediate monocytes (Ly6C^+^MHCII^+^), and mature macrophages (Ly6C^−^ cells) and representative expression of CD11c and CD163 by Ly6C^−^ macrophages. **F.** Representative expression of *Cx3cr1*-GFP, FRý and CD206 expression by CD11c^+^, CD163^+^ and CD11c^-^CD163^-^ macrophage subsets. Data are representative of at least 10 experiments (CD11c, CD163, FRý, CD206) or from 1 experiment CX3CR1-GFP. **G.** Representative immunofluorescence staining of a cross section of distal colon stained for DAPI, CD68 (yellow), CD163 (red) and CD11c (cyan). **H.** Mean number of CD11c/CD163-defined macrophages per 2mm^2^. n=10 C57BL6J naïve male mice from two independent experiments. 3 ROIs quantified per mouse. Symbols represent individual mice. Friedman’s test with Dunn’s post hoc test. Error bars ± SEM. **I.** Immunofluorescence of a cross section of distal colon stained with Hoechst (white) for nuclei, CD31 (red) for blood vessels, Tujj1 (cyan) for nerves and CD163 (yellow). **J.** Fast gene set enrichment analysis (FGSEA) using Hallmark, Reactome and gene ontology biological processes (GOBP) gene set signatures of each macrophage cluster in C vs. the others. Heatmap of leading-edge genes for each pathway, with the associated enrichment plots.

To validate the findings from the scRNAseq analysis, we next performed flow cytometry on dissociated colonic tissue, using expression of CD64, Ly6C and MHCII to identify cells of the monocyte/macrophage lineage and visualise the monocyte ‘waterfall’[8], [9], [10] (**Fig. 1E**). Monocytes were defined as Ly6C^+^MHCII^−/+^ whereas mature macrophages expressed high levels of MHCII and lacked Ly6C expression. All macrophage subsets expressed high levels of CX3CR1, as determined by assessing their expression of green fluorescent protein (GFP) in *Cx3cr1*^+/gfp^ knock-in mice[32] (**Fig. 1E-F**). Within the Ly6C^−^MHCII^+^ compartment, CD11c and CD163 defined three subpopulations of macrophages that were roughly equal in abundance (**Fig. 1E**). Consistent with our scRNA-seq data, CD163^+^ macrophages expressed high and uniform levels of folate receptor (FRβ) and CD206 (**Fig. 1F**). Interestingly, at the protein level, we could find little or no expression of CD206 by CD11c^+^ and CD11c^−^CD163^−^ macrophages. Given the common use of CD4 and Tim4 to define intestinal macrophages[12], [20], we assessed expression of these markers across our CD11c/CD163-defined subsets. While we found ∼60% of CD163^+^ macrophages to express Tim4, a fraction of both CD11c^+^ macrophages and those lacking CD11c and CD163 also expressed Tim4, suggesting that Tim4 is not restricted to a transcriptionally defined subset. Likewise, CD4 expression was present across most subsets, albeit to varying degrees (**Supplementary** Fig. 1F).

Next, we used immunofluorescence to examine the anatomical positioning of our defined macrophage subsets. CD163 expression identified macrophages in the lower mucosa, submucosa and muscularis in both the colon and small intestine, whereas CD11c predominantly defined mucosal macrophages (**Fig. 1G, Supplementary** Fig. 1G. Quantification of macrophage subsets via immunofluorescence staining showed that CD163^+^ macrophages were the most abundant (**Fig. 1H**), a finding that was at odds with our flow cytometry data, suggesting that isolation of these cells by enzymatic dissociation may underestimate their abundance. Further immunofluorescence staining, including markers for endothelial cells (CD31) and nerves (Tuj1), demonstrated an intimate association of CD163^+^ macrophages with blood vessels and nerves, structures which largely parallel each other in the intestine, whereas CD11c^+^ were predominantly subepithelial (**Fig. 1G, I**). These results are consistent with recent characterisation of macrophage subsets in human intestine, where ACP5 (which is coincident with CD11c expression in our mouse dataset) and CD163-defined macrophages are observed in the subepithelial mucosa and deeper mucosa/submucosa, respectively[6], [33].

Given the distinct positioning of these macrophage subsets, we performed Fast Gene Set Enrichment Analysis (FGSEA) to infer their function. *Itgax*^+^ macrophages (Cluster 0) were enriched for genes encoding multiple matrix metalloproteinases (MMPs) (**Fig. 1C, D**), indicative of tissue remodelling capabilities[34]. They were also enriched for genes associated with ‘interleukin 10 signalling’, including the anti-inflammatory cytokine *Il10* itself, suggesting they may align with the previously described population[35], but also the pro-inflammatory mediators *Il1b* and *Il1a* (**Fig. 1J**). In addition, chemokines such as *Cxcl1*, *Cxcl2*, *Cxcl10* and *Ccl4* were all more highly expressed by *Itgax*^+^ macrophages relative to the other macrophage clusters. Cluster 1, the *Cd163^+^*cluster, was found to have gene modules associated with the complement cascade and phagocytosis machinery upregulated. These included *Cd55*, *C2*, *C4a* and *C4b*; and *Fcn1*, *Fcn2* and *Cd36* respectively. Cluster 2 was enriched for ‘Cell surface interactions at the vascular wall’ including *Fn1*, which encodes fibronectin, a glycoprotein that has been shown to be a key component of the extracellular matrix and may play a role in facilitating macrophage-fibroblast interaction[36], [37]. However, another pathway highlighted was ‘Eukaryotic translation initiation’, which can be associated with cells in active/ongoing differentiation[38], suggesting that Cluster 2 may, in part, represent monocyte/macrophage intermediaries. Taken together, these data show that expression of CD11c and CD163 offers a robust approach to define transcriptionally and anatomically distinct intestinal macrophage subsets in mice.

### CD11c/CD163-defined macrophages have distinct replenishment kinetics

Having established robust approaches for defining distinct macrophage subsets, we next sought to evaluate the replenishment kinetics of these subsets using a variety of lineage tracing tools. First, we used tissue-protected bone marrow chimeric mice to demonstrate differences in replenishment across macrophage subsets. Specifically, we found high rates of replenishment amongst CD11c^+^ and CD11c^−^CD163^−^ macrophages, whereas those expressing CD163 had significantly lower levels of replenishment over the same time frame (**Fig. 2A, Supplementary** Fig. 2A). Next, we used *Ms4a3*-based fate mapping, in which all granulocyte macrophage progenitors (GMPs) and their progeny, including monocytes, can be labelled[39]. Thus, labelling of macrophages in this system is indicative of derivation from blood monocytes, whereas lack of label may reflect derivation from embryonic sources. In *Ms4a3*^Cre/+^.*Rosa26*^CAG-LSL-tdTomato/+^ mice (which were also *Cx3cr1*^+/gfp^) Ly6C^hi^MHCII^−^ (‘P1’) monocytes showed similar labelling to their counterparts in blood (**Fig. 2B, Supplementary** Fig. 2B, C), consistent with our previous data showing that these represent newly extravasated monocytes[9]. Interestingly, all subsets of mature macrophages were labelled at levels markedly lower than that of blood monocytes at 3 weeks of age (all <20%), suggesting at this age the majority of colonic macrophages are not derived from haematopoietic stem cell (HSC)-derived monocytes. However, by 12 weeks of age there was a marked increase in tdTomato labelling, consistent with post-natal contribution of HSC-derived monocytes to these compartments[8], [17]. Importantly, there were differences in labelling between the subsets, with the frequency of tdTomato^+^ cells amongst CD11c^+^ macrophages and CD11c^−^CD163^−^ macrophages increasing more quickly with age than within the CD163^+^ compartment. However, by 1 year of age, we did not detect any difference in labelling in any subset compared to that at 12 weeks (**Fig. 2B, Supplementary** Fig. 2C), suggesting that either (1) the rate of monocyte contribution plateaus during adulthood, (2) a fraction of the adult macrophage compartment remains of embryonic origin, or (3) that a fraction of the macrophage compartment is replenished by cells not targeted in the *Ms4a3*^Cre^ system. To test the possibility that a fraction of each subset may represent long-lived macrophages derived from embryonic sources, we made use of *Cdh5*^Cre-ERT2/+^.*Rosa26*^CAG-LSL-tdTomato/+^ mice, in which yolk sac-derived or HSC-derived progenitors (and their progeny) can be labelled through timed administration of 4-hydroxytamoxifen (4-OHT)[40] at either embryonic day 7.5 (E7.5; yolk sac) or E10.5 (HSC), respectively. Delivery of 4-OHT at E7.5 led to high labelling of brain microglia (data not shown), consistent with their yolk sac origin, whereas there was negligible labelling of blood monocytes in these same mice when assessed at 7-11 weeks or 24 weeks (**Fig. 2C, Supplementary** Fig. 2D, J). Within the colon at 7-11 weeks, there was little or no labelling in monocytes, or CD11c^−^CD163^−^ macrophages with CD11c^+^ macrophages not having significantly higher labelling than the monocytes. However, a small but significant proportion (<3%) of CD163^+^ macrophages were labelled (**Fig. 2C, Supplementary** Fig. 2D), suggesting a proportion of these cells remain of embryonic origin at this timepoint. By 24 weeks of age, the frequency of tdTomato labelled CD163^+^ macrophages still had ∼3% labelling, suggesting a very small proportion of CD163^+^ macrophages in the adult colon remain of embryonic origin; a finding consistent with *Cx3cr1*-based fate mapping[4]. We next administered 4-OHT at E10.5 to label HSC-derived cells[40]. In line with this, brain microglia were spared from labelling, whereas colonic Ly6C^+^MHCII^−^ monocytes were fully labelled (**Fig. 2D, Supplementary** Fig. 2E). CD11c^+^ macrophages matched labelling seen in monocytes, whereas CD11c^−^CD163^−^ and CD163^+^ macrophages were not labelled to the same extent as monocytes, consistent with a small but detectable fraction of these cells coming from non-HSC sources and indicating that the CD11c^−^CD163^−^ macrophages may be more heterogeneous and not simply represent monocyte/macrophage intermediaries.

**Figure 2.**
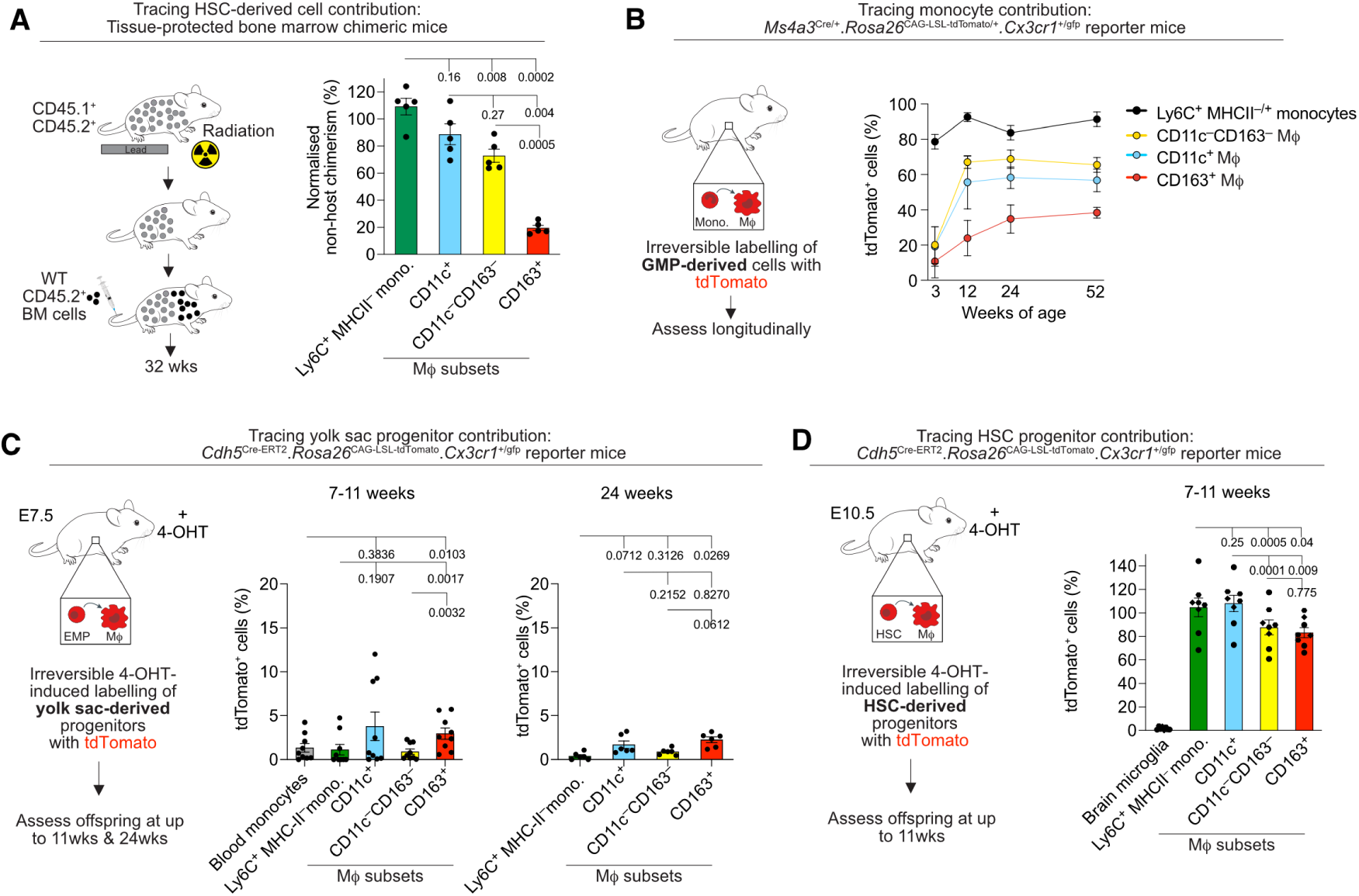
Colonic macrophage subsets have distinct replenishment kinetics. **A.** Scheme for the generation of tissue-protected BM chimeric mice and normalised non-host chimerism of Ly6C^+^MHCII^-^ monocytes and CD11c/CD163-defined macrophages from tissue-protected BM chimeric mice 36 weeks post-reconstitution. Data from one experiment. n=5 mice. All mice used were male. Paired one-way ANOVA with Tukey’s post hoc test. Error bars ± SEM. **B.** Experimental scheme of *Ms4a3*^Cre/+^.*Rosa26*^CAG-LSL-tdTomato^ lineage-tracing mice to trace the progeny of GMPs and mean frequency of tdTomato^+^ of Ly6C^+^MHCII^-/+^monocytes and CD11c/CD163-defined macrophages at 3, 12, 24 and 52 weeks of age. Data from a single experiment (24 and 52 weeks of age) with n=3-4 mice per group or pooled from 2 individual experiments (3 and 12 weeks of age) with n=9 mice per group. Both male and female mice were used and analysed together. Error bars ± SD. Mixed effects analysis with Šídák’s multiple comparisons test. **C.** Experimental scheme of *Cdh5*^Cre-ERT2/+^.*Rosa26*^CAG-LSL-tdTomato/+^.*Cx3cr1*^+/GFP^ fate mapping mice and mean frequency of tdTomato labelled Ly6C^+^MHCII^-^ colonic monocytes and CD11c/CD163-defined macrophages from 7-11 week old (*left*) or 24 week old (*right*) *Cdh5*^Cre-ERT2/+^.*Rosa26*^CAG-LSL-tdTomato/+^.*Cx3cr1*^+/GFP^ mice. Labelling is normalised to tdTomato labelling of brain microglia. Data are pooled from three independent experiments with n=9 mice (7-11 weeks) or from n=6 (24 weeks). Both male and female mice were used and analysed together. Paired one-way ANOVA with Tukey’s multiple comparisons test. **D.** Experimental scheme of *Cdh5*^Cre-ERT2/+^.*Rosa26*^CAG-LSL-tdTomato/+^.*Cx3cr1*^+/GFP^ fate mapping mice and mean frequency of tdTomato labelled brain microglia, Ly6C^+^MHCII^-^ colonic monocytes and CD11c/CD163-defined macrophages from 7-11 week old *Cdh5*^Cre-ERT2/+^.*Rosa26*^CAG-LSL-tdTomato/+^.*Cx3cr1*^+/GFP^ mice with labelling normalised to tdTomato labelling of classical Ly6C^hi^ blood monocytes. Data are pooled from three independent experiments with n=8 mice (7-11 weeks). Both male and female mice were used and analysed together. Paired one-way ANOVA with Tukey’s multiple comparisons test. Symbols represent individual mice.

Taken together, these data show that embryonic progenitors contribute to all colonic macrophage subsets, but that these are largely displaced by HSC-derived cells in the post-natal period, consistent with previous work[8], [17]. However, the rate of replenishment of colonic macrophages is not uniform, with sub-epithelial CD11c^+^ macrophages replenished more quickly than CD163^+^ macrophages in the lower mucosa, sub-mucosa and muscularis. Finally, given the paucity of embryonic-derived macrophages in the adult colon with incomplete labelling of these cells in *Ms4a3*^Cre/+^.*Rosa26*^CAG-LSL-tdTomato/+^ mice, our findings indicate that progenitors that are not captured by the *Ms4a3*^Cre^ system likely contribute to the replenishment of colonic macrophages in steady state.

### Intestinal macrophages accumulate in a model of resolving colitis

Next, to understand how inflammation impacts the macrophage compartment during injury and its subsequent resolution, we used a model of self-resolving colitis by feeding mice with 2% dextran sodium sulphate (DSS) for four days before allowing them to recover for two weeks (day 18). Here, we observed peak inflammation at day 3-4 after cessation of DSS, as evidenced by the weight loss, clinical score (based on general appearance, faecal consistency and evidence of blood in the stool), colon shortening and granulocyte accumulation (**Fig. 3A-C**). However, by day 18, each of these indices had returned to baseline levels, including colon architecture (**Fig. 3A-C, G**). DSS administration led to marked expansion of Ly6C^hi^MHCII^−^ and Ly6C^+^MHCII^+^ monocytes at day 8 and these had returned to normal numbers by day 18 (**Fig. 3D-F**). The total number of Ly6C^−^ macrophages also increased at this time before returning to baseline by day 18 (**Fig. 3D-F**). Flow cytometry analysis of the macrophage subsets indicated that the increase in macrophages reflected increased abundance of CD11c^+^ and CD11c^−^ CD163^−^ macrophages, with no change to the CD163^+^ compartment (in this experiment defined on the basis of FRβ expression) (**Supplementary** Fig. 3A). However, complementary immunofluorescence imaging revealed that all macrophage subsets significantly increased in number during acute colitis, with the CD11c^−^CD163^−^ and CD11c^+^ macrophage numbers elevated up until day 11 (**Fig. 3G, H**). Moreover, each subset could be identified in areas of severe damage, including disruption to the crypt architecture (**Fig. 3G**). The numbers of CD11c^−^ CD163^−^ and CD163^+^ macrophages returned to baseline by day 18, while CD11c^+^ macrophages remained elevated at day 18 during resolution (**Fig. 3H**). Thus, in contrast to many published studies describing a loss of mature, resident type macrophages in this model[9], [11], [41], [42], our imaging data show that mature macrophages appear to remain during inflammation and are present in inflammatory lesions.

**Figure 3.**
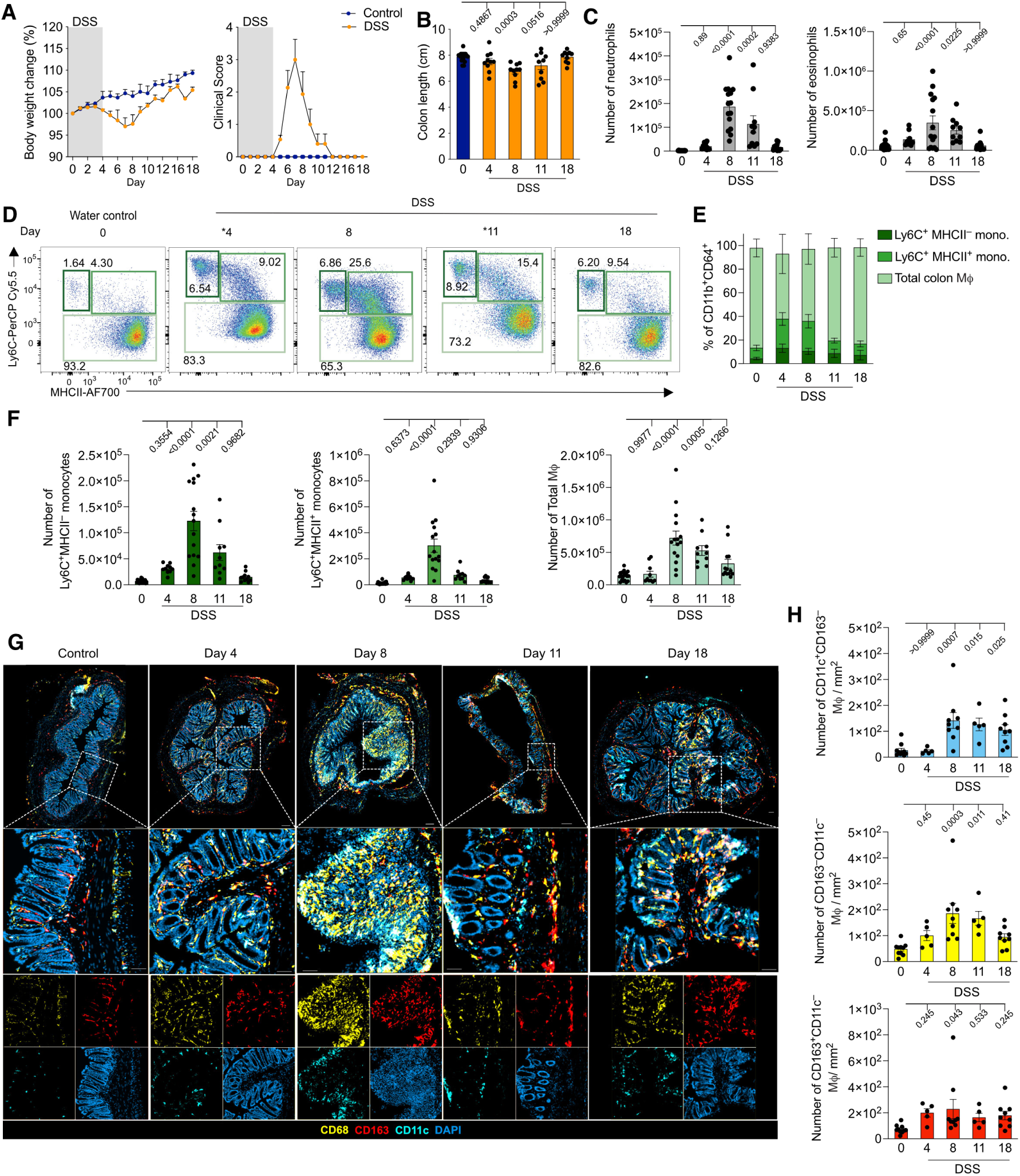
Subset-specific changes to macrophages during and following colitis. **A-B.** Weight loss and clinical score (**A**) and colon length (**B**) of male mice fed 2% DSS for four days and returned to normal drinking water for up to fourteen days (‘recovery colitis’). Error bars ± SD. One-way ANOVA with Dunnett’s multiple comparisons test. **C**. Absolute number of eosinophils (*left*) or neutrophils (*right*) from male control mice (0) or male mice harvested at the indicated timepoints of the recovery colitis model. Data from three independent experiments with n=21 (control mice, day 0), n=15 (day 8), n=14 (day 18), or from two independent experiments with n=10 mice (day 4 and 11). Mice which reached humane endpoints were culled prior to their respective experimental day and were excluded from the experiment. One-way ANOVA with Dunnett’s multiple comparisons test. Error bars ± SEM. **D.** Representative expression of Ly6C and MHCII by Lin^−^CD11b^+^CD64^+^ leukocytes to identify recently infiltrated Ly6C^+^MHCII^−^ monocytes; intermediate Ly6C^+^MHCII^+^ monocytes; and Ly6C^−^MHCII^+^ macrophages across the recovery colitis model. * Represents flow experiments performed on days distinct from other representative plots. **E-F.** Mean frequency **(E)** and number **(F)** of Ly6C^+^MHCII^−^ monocytes, Ly6C^+^MHCII^−^ monocytes and total Ly6C^−^ macrophages across the recovery colitis model from mice in **C**. **G-H.** Representative immunofluorescence staining (**G**) of a cross section of distal colon taken at the indicated time points of the recovery colitis model stained for CD68 (yellow), CD163 (red) and CD11c (cyan) and DAPI (blue). Increased magnification image highlighted by a dotted white line is a representative 250mm^2^ region of interest (ROI). Mean number of CD11c^+^ (CD11c^+^CD68^+^CD163^-^), CD11c^-^CD163^-^M< (CD11c^-^CD163^-^CD68^+^) and CD163^+^ (CD163^+^CD68^+^CD11c^-^) macrophages per 250mm^2^ (**H**). Data are from n=10 (controls, D0), n=5 (day 4, 11) or n=9 (day 8 and 18) pooled from two independent experiments. 3 ROIs (250mm^2^) based on DAPI staining were quantified per mouse. 12μm thick sections. One-way ANOVA with Dunnett’s post hoc test. Error bars ± SEM. Symbols represent individual mice.

### Macrophages in the post-inflammation colon are transcriptionally distinct to their homeostatic counterparts

Previous work using bulk transcriptional analysis has suggested that mature colonic macrophages show very few transcriptional changes during acute colitis[11]. However, if and how specific macrophage subsets change once inflammation has resolved has not been assessed. While our immunofluorescence results above suggest that the mature macrophage compartment expands in DSS colitis, whether these cells reflect homeostatic macrophages that persist through inflammation into resolution, or the progeny of recruited monocytes elicited during inflammation, or indeed both, remained unclear. To address this, we used another genetic fate mapping model which exploits CX3CR1 expression by homeostatic colonic macrophages. In *Cx3cr1*^Cre-ERT2/+^.*Rosa26*^LSL-RFP/+^ mice, tamoxifen administration leads to irreversible labelling of *Cx3cr1*-expressing macrophages with red fluorescent protein (RFP) (**Fig. 4A**). We established an optimal tamoxifen dose to label colonic macrophages to high efficiency, using CX3CR1^hi^ microglia as a positive control (**Supplementary** Fig. 3B-H). Next, we used these *Cx3cr1*^Cre-ERT2/+^.*Rosa26*^LSL-RFP/+^ mice in our recovery colitis model and determined if, and to what extent, induction of colitis altered the number of RFP-labelled macrophages at recovery (**Fig. 4B**). Importantly, we built in a one week ‘wash out’ period following tamoxifen administration (and prior to DSS administration) to ensure the small levels of RFP labelling within (short-lived) blood Ly6C^hi^ monocytes were no longer present (**Supplementary** Fig. 3I, J), since their recruitment could have confounded our interpretation of RFP^+^ cell persistence/accumulation. At baseline, a high level of RFP-labelling was detected across the macrophage subsets by flow cytometry (**Supplementary** Fig. 3K-N), however, to ensure we had sufficient representation of all macrophage subsets, we opted to use image analysis to determine the dynamics of the RFP labelled cells. Using this approach, we found that in the lamina propria ∼40% of CD11c^+^ and CD163^+^ macrophages were labelled in control *Cx3cr1*^Cre-ERT2/+^.*Rosa26*^LSL-RFP/+^ mice (administered tamoxifen but not DSS) three weeks after the final dose (**Fig. 4C**). We found that the labelling of CD11c^−^CD163^−^ macrophages was higher at ∼75%, although the basis for this is unclear. Importantly, we found that induction of colitis did not alter the frequency of RFP^+^ cells in any of the macrophage populations (**Fig. 4C**), suggesting that the macrophages labelled with RFP prior to colitis induction persist through inflammation into the recovery phase. Importantly, during peak colitis (day 8) RFP^+^ macrophages could be identified throughout the mucosa, including in neutrophil-rich, inflamed areas, which lacked epithelial integrity, evident by a loss of EpCAM (**Supplementary** Fig. 4A). Next, to understand if periods of inflammation alter colonic macrophages and, in particular, change the nature of previously homeostatic macrophages, we combined our *Cx3cr1*-based fate mapping with scRNAseq. Live colonic CD45^+^ CD11b/CD11c^+^ non-granulocytes from *Cx3cr1*^Cre-ERT2/+^.*Rosa26*^LSL-RFP/+^ mice at day 18 of the recovery colitis model, or controls (no DSS) were isolated and sequenced using the 10X platform. Again, cells of the monocyte/macrophage lineage were identified on the basis of their expression of *Csf1r, Adgre1, Fcgr1, Cx3cr1* and lack of cDC lineage genes *Itgae, Dpp4, and Xcr1;* or contaminating B cell genes i.e. *Cd19* (**Supplementary** Fig. 4B,C). Again, for increased granularity of the macrophage populations, monocytes were removed on the basis of their *Ly6c2* expression (**Supplementary** Fig. 4D,E). Broadly similar clusters were identified as in the naïve mice (**Fig. 4D, E; Supplementary** Fig. 4F, G). Cluster 0 represented the major *Itgax* expressing cluster and was defined by *Col14a1*, *Col15a1* and *Lilra5* expression (**Supplementary** Fig. 4G). Cluster 1 was also *an Itgax-*expressing cluster, but was defined by higher expression of *Pgf, Vegfa, and Nrg1.* Cluster 2, the *Cd163^+^* cluster, was again defined by *Folr2, Lyve1*. Cluster 3 appeared to align with the CD11c^−^CD163^−^ population. It was enriched in genes such as *Ccr2*, *Plac8, S100a8* and *S100a6*, which are typically associated with monocytic cells, suggesting these might represent cells recently derived from monocytes. An additional small cluster (4) was defined by *Tspan10, and Lrp8* (**Supplementary** Fig. 4G). The rarity of this cluster did not permit meaningful analysis and so was not examined further.

**Figure 4.**
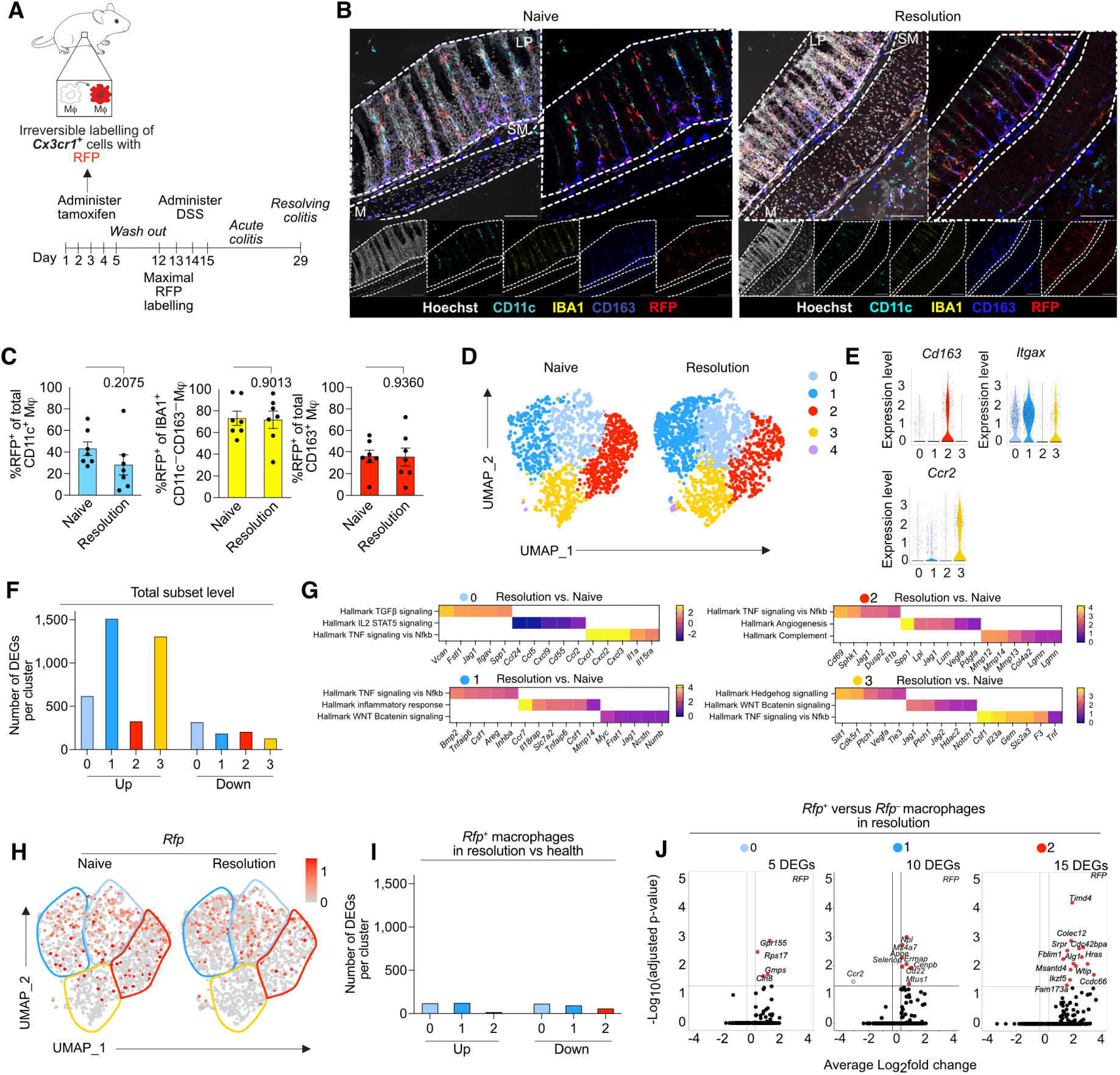
Homeostatic macrophages persist through inflammation and adapt to aid repair. **A.** Experimental scheme for *Cx3cr1*^Cre-ERT2^.*Rosa26*^LSL-RFP/+^ fate-mapping **B.** Immunofluorescence staining of Hoechst (grey), CD11c (cyan), IBA1 (yellow), CD163 (blue) and RFP (Red; representing RFP^+^ fate-mapped macrophages) in tamoxifen treated *Cx3cr1*^Cre-ERT2/+^.*Rosa26*^LSL-RFP/+^ mice that had received a 4 day course of DSS 18 days earlier or non-DSS treated controls. 12μm thick Swiss roll sections. Data are from n=7 naive and n=7 in resolution. LP-Lamina propria, SM-muscularis, M-muscularis. **C.** Quantification of the proportion of RFP^+^ macrophage subsets normalised to area, from immunofluorescence imaging. Data are from two repeat experiments mixed male and female. N=7. **D.** UMAP projection of macrophages (defined as *C1qa*^+^*Ly6c2^−^*)) obtained from *Cx3cr1*^Cre-ERT2/+^.*Rosa26*^LSL-RFP/+^ mice pulsed with tamoxifen, rested for 1 week and then fed DSS for 4 days before recovering for 14 days, or tamoxifen-treated control mice maintained on normal drinking water. Violin plots of cluster defining genes. **E.** Violin plots showing the cluster defining genes for clusters defined in **D**. **F.** Number of differentially expressed genes (DEGs) between DSS treated and controls per cluster. **G.** Fast gene set enrichment analysis (FGSEA) using Hallmark gene set signatures of each cluster in resolution vs naïve. Heatmap of leading-edge genes for each pathway. **H.** *Rfp* expression across clusters in **D**. in resolution and naive mice. **I.** Number of differentially expressed genes (DEGs) between DSS treated and controls per cluster focussed on *Rfp*^+^ cells. **J.** Volcano plots showing differentially expressed genes (DEGs) by the *Rfp^+^* cells compared with *Rfp*^−^ cells.

First, we examined if CD11c/CD163-defined macrophages in the post-inflammation colon differed to those found in naïve mice. In resolution, there were 929 differentially expressed genes (DEGs) in Cluster 0 (*Itgax^+^Itga6^+^*) compared with equivalent cells in health (**Fig. 4F**). There were over 1692 DEGs between resolution and health in the additional *Itgax*^+^*Nrg1*^+^ population (Cluster 1), perhaps explaining its separation from the main *Itgax* expressing cluster. There were fewer changes in the longer-lived *Cd163*^+^ cluster, though there were still 527 DEGs in resolution compared with the same cluster in health. Similar to the *Itgax*^+^ cluster, there were marked differences within the *Itgax^−^Cd163*^−^ cluster, with 1428 DEGs in resolution compared with its counterparts in health (**Fig. 4F**).

Gene set enrichment analysis revealed that there was a general enrichment of TGFβ, WNT and Hedgehog signalling (**Fig. 4G, Supplementary 4H**), which is consistent with a tissue repair role through promotion of epithelial stem cells support. For the *Itgax^+^Itga6^+^* cluster, within the module of genes implicated in TGFβ signalling, *Itgav*, which encodes integrin av, was one of the most upregulated (**Supplementary 4I**). Integrin αv can activate latent TGFβ, and *Spp1* (encoding osteopontin), which is implicated in tissue repair[43]. Loss of *Itgav* in myeloid cells leads to development of spontaneous colitis[44], reaffirming a regulatory/repair role. This cluster also had an upregulation of a gene set implicating ‘TNF signalling’ in resolution, but downregulation of modules associated with ‘IL2/STAT5’ signalling (**Fig. 4G**). TNF and inflammatory signalling gene sets, along with WNT signalling, were upregulated by the *Itgax*^+^*Nrg1*^+^ cluster (**Fig. 4G**). *Mmp14* (matrix metalloproteinase 14), implicated in tissue remodelling, and *Vegfa* (vascular endothelial growth factor alpha), important for tissue repair, were both upregulated by this cluster[45] (**Supplementary 3I**). As was *Nrg1* itself, which encodes neuregulin1, and has been found to define a macrophage population which expands in ulcerative colitis and was associated with mucosal healing through cross-talk with epithelial stem cells[46]. The *Cd163*^+^ macrophages had upregulation of gene sets associated with angiogenesis, which are also involved in tissue repair (**Fig. 4G, Supplementary 3I**). Moreover, they displayed upregulated gene sets including genes encoding MMPs (*Mmp12, Mmp13* and *Mmp14*), indicative of a role in tissue remodelling that was not as evident in these cells during health. The *Itgax*^−^*Cd163*^−^ cluster had increased expression of modules associated with repair (e.g., Hedgehog/WNT β catenin signalling) but also with inflammation, with upregulated TNF signalling indicating a duality in their function or further heterogeneity amongst this cluster (**Fig. 4G, Supplementary 3I**). Thus, despite looking phenotypically similar by flow cytometry analysis and positioning by imaging, macrophages in the post-inflammation colon are transcriptionally very different to their counterparts in health.

Finally, we sought to determine if there were changes to the macrophages present before the onset of inflammation that had persisted into the post-inflammation environment, and which are thought to have restricted plasticity. To assess this, we compared both *Rfp*^+^ cells in health with *Rfp*^+^ cells post-inflammation, and *Rfp*^+^ cells post inflammation with their *Rfp*^−^ counterparts. We detected a high number of DEGs between the *Rfp*^+^ cells in health and their counterparts in recovery across all subsets, although notably fewer in the *Cd163*^+^ cluster (**Fig. 4I**). This suggested that, in contrast to previous work suggesting resident macrophages do not change during acute colitis[11], long-lived resident macrophages in the post-inflammatory environment adapt to aid tissue repair. Irrespective of their positioning in the gut wall, all *Rfp*^+^ macrophages in the post-inflammation environment exhibited upregulation of the pathways identified at population level (e.g., TGFβ signalling for the *Itgax* expressing clusters (0 and 1); WNT signalling for the *Itgax*^+^*Itga6*^+^ and the *Cd163*^+^ subsets; and increased Hedgehog and Notch signalling within the *Itgax*^+^*Nrg1*^+^ cluster) (**Supplementary 3J**). Intriguingly, comparison of *Rfp*^+^ cells with *Rfp*^−^ cells in the post-inflammation environment within the same sample showed little transcriptional difference (**Fig. 4J**), suggesting *de novo* recruited (RFP^−^) and long-lived cells (RFP^+^) are transcriptionally similar; and confirming the environment and not the time of residency is the dominant factor in determining the macrophage state in this context.

Taken together, our data show that the colonic macrophage compartment is diverse, with transcriptionally, anatomically and developmentally distinct macrophage subsets in health. Inflammation results in marked changes to the composition of the monocyte/macrophage compartment, and while the homeostatic composition appears to reestablish during resolution, all macrophage subsets adopt a distinct transcriptional and functional state associated with tissue repair. Given many of these pathways become dysregulated in the context of chronic inflammation and fibrosing disease, understanding the factors which govern these pathways could allow novel targeting of macrophages in these contexts. Importantly, understanding whether this leads to an altered response with subsequent inflammatory events remains unexplored and should be examined in future work.

## Material and Methods

### Mice and ethics

All mice used were of a C57BL/6J background. All experiments followed the Animal Research: Reporting of In Vivo Experiments (ARRIVE) guidelines and were approved by the UK Home Office (project licence: PP6581698). Mice were housed under standard specific pathogen-free (SPF) conditions, with unrestricted access to food and water, and maintained on a 12-hour light/dark cycle. Male and female mice were used, with the sexes and ages noted in the figure legends. To normalise microbiota between cages, bedding was mixed across cages for 2-3 weeks prior to experimental start dates. The transgenic mice used in this study are listed in Table 1.

**Table 1.** Mouse strains.

| Strain | Source | Ref./JAX ref. |
| --- | --- | --- |
| C57BL/6J | Charles River Laboratories | N/A |
| C57BL/6J CD45.2 | University of Edinburgh | N/A |
| C57BL/6J CD45.1<sup>+</sup>/CD45.2<sup>+</sup> | University of Edinburgh | N/A |
| *Ms4a3*<sup>Cre</sup>.*Rosa26*<sup>CAG-LSL-tdTomato</sup> | Generated by Prof. F.Ginhoux (ASTAR) and obtained through collaboration with Dr. Rebecca Gentek (CRH, University of Edinburgh) | [39] |
| *Cdh5*<sup>Cre-ERT2</sup>.*Rosa26*<sup>CAG-LSL-tdTomato</sup>.*Cx3cr1*<sup>+/GFP</sup> | Kindly provided by Dr. Rebecca Gentek (CRH, University of Edinburgh) | [40] |
| *Cx3cr1*<sup>GFP/GFP</sup> | Kindly provided by Prof. Jeff Pollard (CRH, University of Edinburgh) | [32] |
| *Cx3cr1*<sup>CreERT2/+</sup>.*Rosa26*<sup>LSL.RFP/+</sup> | Generated by crossing double homozygous *Cx3cr1*<sup>CreERT2/CreERT2</sup>.*Rosa26*<sup>LSL.RFP/LSL.RFP</sup> mice with C57BL/6 mice | [47], [48], [49] |

### Induction of dextran sulfate sodium (DSS) induced colitis

To induce acute colitis, mice received 2% dextran sulfate sodium (DSS) salt (reagent grade; 36,000 - 50,000 kDa; MP Biomedical) *ad libitum* in sterile drinking water for up to 4 days. To induce self-resolving colitis, mice were fed 2% DSS for 4 days and then switched back to normal drinking water for up to 14 days. Mice were checked daily for bodyweight change, rectal bleeding, stool consistency and general appearance with a clinical score generated. Mice which lost >20% of their starting bodyweight were culled immediately as per Home Office guidelines.

### Generation of tissue-protected bone marrow chimeric mice

Bone marrow was flushed from the femurs and tibiae of CD45.2^+^ mice using sterile PBS and physically dissociated using a syringe (without needle) and strained through a 40μm strainer. This cell suspension was then spun down at 300g for 5 mins, the supernatant discarded, before being incubated in 1x RBC lysis buffer (Sigma; 11814389001) for 3 mins at room temperature. The cells were washed in ice cold sterile PBS and cells counted manually using a haemocytometer. 5×10^6^ cells were intravenously transferred into sex-matched CD45.1/.2^+^ mice which had been irradiated with a 9.5 Gy γ-irradiation with lead protecting all but the head and upper torso. Recipient mice were examined for the presence of donor cells 36 weeks after the transfer.

### Induction of Cre in fate mapping models

To induce Cre in *Cx3cr1*^CreERT2/+^.*Rosa26*^LSL.RFP/+^ mice, 3 mg tamoxifen (Sigma) dissolved in sesame oil (Sigma) was administered by oral gavage for 5 consecutive days. For the *Cdh5*^CreERT2/+^.*Rosa26*^CAG-LSL-tdTomato^.*Cx3cr1*^gfp/+^ system, female mice aged 6–10 weeks were timed-mated, and the presence of vaginal plugs the morning after mating was considered embryonic day 0.5 (E0.5). To induce reporter recombination in the developing offspring, a single dose of 4-hydroxytamoxifen (4-OHT, 1.2 mg) (Sigma-Aldrich) was administered to pregnant females via IP injection at E7.5 or E10.5. To mitigate the adverse effects of 4-OHT on pregnancy, progesterone (0.6 mg) (Sigma-Aldrich) was co-administered in corn oil. In cases where females were unable to deliver naturally, pups were delivered via cesarean section and cross-fostered with CD1 lactating mice.

### Cell Dissociation

Colonic leukocytes were isolated from enzymatically digested tissue as previously described[50]. Briefly, colons were excised and soaked in RPMI/2% FCS before being opened longitudinally, rinsed in the RPMI/2% FCS, and cut into 0.5cm pieces. To remove epithelial cells, tissue was incubated with calcium and magnesium-free (CMF) Hank’s Balanced Salt Solution (HBSS; Gibco) containing 2mM EDTA (Gibco) in a shaking incubator 37°C for 15 minutes. Samples were then shaken vigorously and the supernatant discarded by passing through nitex mesh. Tissue pieces were collected, and this step was repeated for a further 15 minutes in fresh CMF HBSS/2mM EDTA. Following the second incubation the tissue was rinsed with pre-warmed CMF HBSS without EDTA. The tissue was digested in 5 ml pre-warmed RPMI/10% FCS containing 0.425mg/ml Collagenase V (Sigma, C9263-1G), 0.625mg/ml Collagenase D (Roche, 11088882001), 1mg/ml Dispase (Gibco, 17105-041), 30μg/ml DNase (Roche,101104159001) for 30-35 minutes in a shaking incubator at 37°C. To increase digestion, the falcon tube was shaken vigorously every 5-7 minutes. 25 ml ice cold FACS buffer (PBS/2mM EDTA/2% FCS) was then added to the samples to stop the enzymatic reaction, before the cell suspension was passed through a 40μm cell strainer (Fisher Scientific). Cells were spun at 400g for 5 minutes at 4°C, supernatant discarded and resuspended in ice cold FACS buffer (PBS/2mM EDTA/2% FCS) and kept on ice until further use.

To isolate microglia, brains were removed from terminally anaesthetised mice and stored in RPMI/2% FCS. Then, brains were chopped into small pieces with a razor blade, before being digested with the enzyme mix used for colonic tissue (5ml/brain) for 40 minutes in a shaking incubator at 37°C. 25 ml ice cold FACS buffer was then added to the samples to stop the enzymatic reaction, before the cell suspension was passed through a 100μm cell strainer (Fisher Scientific).

### Processing of whole blood for flow cytometry

Where blood leukocytes were examined, mice were culled by overdose with an intraperitoneal injection of sodium pentobarbital. Needles flushed with 0.5M EDTA (Invitrogen) were used to draw 0.1-0.2 ml blood from the inferior vena cava. To lyse red blood cells, for every 0.1 mL of blood 0.9 ml of 1x red blood cell (RBC) lysis buffer (Biolegend) was added, vortexed and incubated on ice for 5 mins. Cells were washed in 9 ml ice-cold FACS buffer and spun down at 300g for 5 minutes at 4°C and the supernatant discarded. This was then repeated, and the cells were resuspended in FACS buffer and kept on ice until use.

### Flow cytometry

1.5-3×10^6^ cells were incubated with anti-CD16/32 (Biolegend) to reduce non-specific binding of Fc receptors. Cells were then stained with appropriate antibodies (Table 2) at 4°C in the dark for 20-30 minutes. Cells were washed once in FACS buffer and then, if necessary, incubated with fluorochrome-conjugated streptavidin for 15 minutes on ice. Cells were again washed in FACS buffer (PBS/2mM EDTA/2% FCS) and analysed on the 6-laser LSR Fortessa (BD Biosciences). 15 μl of 7-aminoactinomycin D (7-AAD; Biolegend) was added to each sample immediately before acquisition to allow exclusion of dead cells. All flow cytometry data generated was analysed with FlowJo software v10 (BD Bioscience).

| **Antibody** | **Clone** | **Fluorophore** | **Catalogue #** | **Supplier** | **Conc.** |
| --- | --- | --- | --- | --- | --- |
| CCR2 | REA538 | biotin | 130-128-939 | Miltenyi | 1in100 |
| CD115 | AFS98 | APC | 135510 | Biolegend | 1in200 |
| CD11b | M1/70 | APC/ Fire 750 | 101262 | Biolegend | 1in200 |
| CD11b | M1/70 | PECy7 | 101216 | Biolegend | 1in200 |
| CD11c | N418 | BV605 | 117334 | Biolegend | 1in100 |
| CD11c | N418 | BV785 | 117336 | Biolegend | 1in100 |
| CD163 | S15049I | APC | 155306 | Biolegend | 1in400 |
| CD163 | S15049F | PE | 156704 | Biolegend | 1in400 |
| CD163 | S15049I | PECy7 | 155320 | Biolegend | 1in400 |
| CD169 | 3D6.112 | PE–Dazzle | 142424 | Biolegend | 1in200 |
| CD19 | eBio1D3 | PECy5 | 2410805 | Invitrogen | 1in200 |
| CD19 | 6D5 | biotin | 115504 | Biolegend | 1in200 |
| CD206 |  | biotin | MCA2235B | BIORAD | 1in200 |
| CD3 | 17A2 | biotin | 100244 | Biolegend | 1in200 |
| CD3e | 145-2C11 | PECy5 | C0031102920553 | Tonbo Biosciences | 1in200 |
| CD45 | 30-F11 | BV510 | 103138 | Biolegend | 1in200 |
| CD45.1 | A20 | AF488 | 110706 | Biolegend | 1in200 |
| CD45.2 | M5/114.15.2 | AF700 | 107622 | Biolegend | 1in200 |
| CD64 | X54-5/7.1 | BV421 | 139309 | Biolegend | 1in200 |
| CX3CR1 | SA011F11 | APC | 149007 | Biolegend | 1in800 |
| F4/80 | BM8 | PECy7 | 123114 | Biolegend | 1in200 |
| FOLRβ | 10/FR2 | APC | 153306 | Biolegend | 1in200 |
| Ly6C | HK1.4 | PerCP–Cy5.5 | 128012 | Biolegend | 1in200 |
| Ly6C | HK1.4 | APC | 128028 | Biolegend | 1in200 |
| Ly6G | 1A8 | AF488 | 127625 | Biolegend | 1in200 |
| Ly6G | 1A8 | PECy5 | E-AB-F1108G | Elabscience | 1in200 |
| Ly6G | 1A8 | biotin | 127606 | Biolegend | 1in200 |
| NK1.1 | PK136 | PECy5 | 108716 | Biolegend | 1in200 |
| NK1.1 | PK136 | biotin | 108704 | Biolegend | 1in200 |
| Rat MHCII | M5/114.15.2 | AF700 | 107622 | Biolegend | 1in200 |
| SiglecF | REA798 | biotin | 130-112-171 | Miltenyi | 1in200 |
| SiglecF | REA798 | FITC | 130-112-178 | Miltenyi | 1in50 |
| Streptavidin | N/A | BV605 | 405229 | Biolegend | 1in500 |
| Streptavidin | N/A | BV650 | 405231 | Biolegend | 1in500 |
| TIM4 | RMT4-54 | PECy7 | 130010 | Biolegend | 1in200 |
| TruStain FcX | S17011E |  | 156603 | Biolegend | 1in1000 |

### Transcriptional profiling by scRNAseq

#### Isolation of mononuclear phagocytes by fluorescence activated cell sorting

Male mice were euthanised and colons were processed as described for flow cytometry above. Cells were sorted using a FACS Aria II (BD Bioscience). Colonic myeloid cell subsets were sorted by excluding DAPI, CD3, CD19, Ly6G, SiglecF and NK1.1 positive cells amongst CD45^+^ cells and selecting cells expressing either CD11c or CD11b (full gating shown in **Supplementary** Figure 1A). Cells were sorted into PBS/10% FCS, counted, washed in PBS/10% FCS and then processed using the Chromium Single Cell Platform using the 10X Chromium Next GEM version 3.1 Single Cell 3’ Library and Gel Bead Kit, (10X Genomics) and the 10X chromium single cell G Chip kit (10X Genomics) as per the manufacturer’s instructions. Cells were loaded onto the 10X Chromium chip where the cells are emulsified with 10X barcoded gel beads in an oil capsule. This process causes cell lysis, barcoded RNA reverse transcription, amplification, fragmentation, and attachment of a 5’ adaptor and sample index. Libraries were sequenced on the Illumina HiSeq platform (Illumina, Inc) by Genewiz (Azenta Life Sciences). Generation of libraries was performed in house.

#### scRNAseq pre-processing

Pre-processing of the data was performed using R-studio and the Seurat package v4.0 following the Satija lab pipeline[51]. Ambient RNA was removed by comparing raw and filtered matrices with the SoupX package[52] determining the level of contaminating RNA. Adjusted matrices were then analysed using Seurat. Samples were normalised with the SCTransform function, and cells with more than 10% mitochondrial reads were excluded. Likely doublets were removed with the DoubletFinder package[53]. When merging samples the Harmony package[54] was used to correct for batch effects. Variable genes, cell clustering, UMAP and heatmap visualisations were all performed using the Seurat package. Genes used to identify subpopulations were found with the ‘findMarker’ function in Seurat. Volcano plots were created with the ggplot package.

### Preparation of tissue for immunofluorescence

For preparation of ‘Swiss rolls’, colons were excised, soaked in PBS, opened longitudinally and washed in fresh ice-cold PBS. Colon tissue was placed on tissue paper soaked with ice cold PBS, mucosal surface facing up. To preserve crypt architecture, antigen fix (Diapath) was dropped onto the surface before being rolled from the distal end by pushing a 0.1cm piece of the distal end towards the proximal end using a curved forceps. ‘Swiss rolls’ were then fixed in antigen fix for 1 hour on ice. Alternatively, a 0.3cm piece of distal colon was cut off and fixed in antigen fix (Diapath) for 1 hour on ice. The Swiss rolls or pieces of distal colon were then washed in PBS, once quickly and then twice for ten minutes, before being placed in 34% sucrose solution in the fridge overnight. Tissue was removed from the sucrose solution with the excess patted away with white roll. The lumens of the distal colons were carefully filled with 50:50 OCT:PBS solution to preserve structures before being snap frozen in OCT and stored at −80°C. 12μm cross sections were cryosectioned (Leica) and stored at −80° C until use.

### Immunofluorescence staining

Sections were brought to room temperature, outlined with hydrophobic pen (Vector) and washed in PBS/0.2% BSA (Sigma Aldrich). Sections were blocked for 20 minutes with 0.2% CD16/32 (Biolegend) in PBS/1% BSA (Sigma Aldrich), or 0.5% saponin if permeabilising the tissue. Sections were incubated with conjugated or unconjugated primary antibodies or isotype controls (**Table 3**) in humidified chambers for 1 hour or overnight at room temperature. Sections were washed three times in PBS 0.2% BSA (Sigma Aldrich). Secondary antibodies were added to the sections for 1 hour at room temperature. Sections were then washed and, if necessary, incubated with further antibodies before staining nuclei with DAPI, washing and mounting slides with coverslips using Fluoromount-G mounting medium (Invitrogen).

**Table 3.**
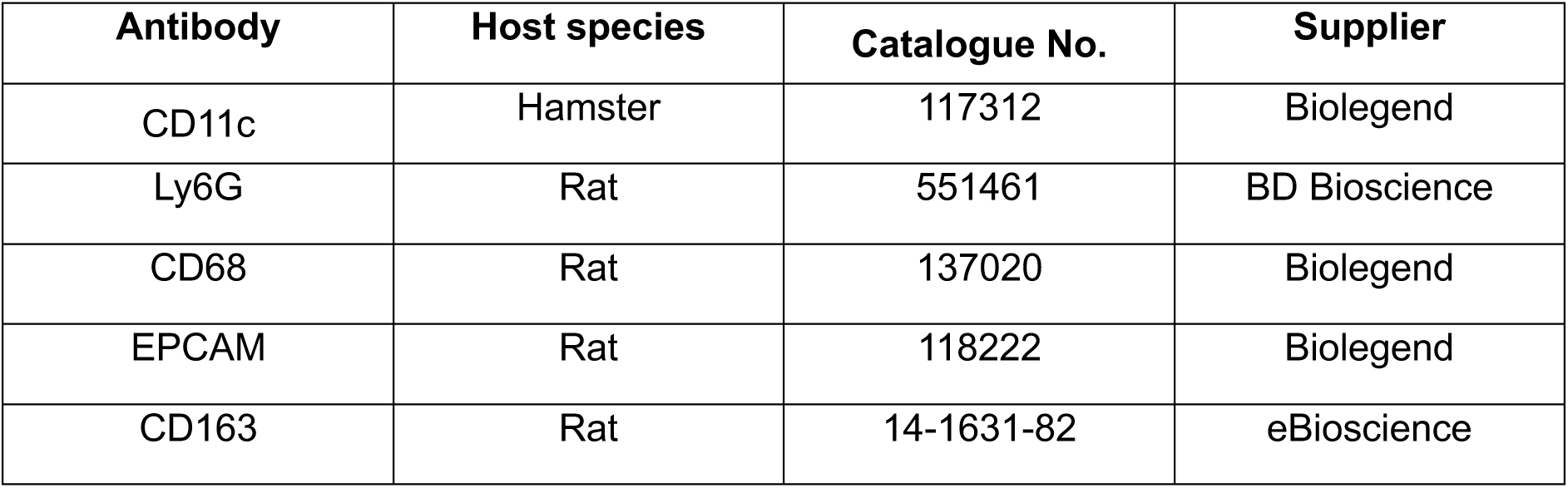
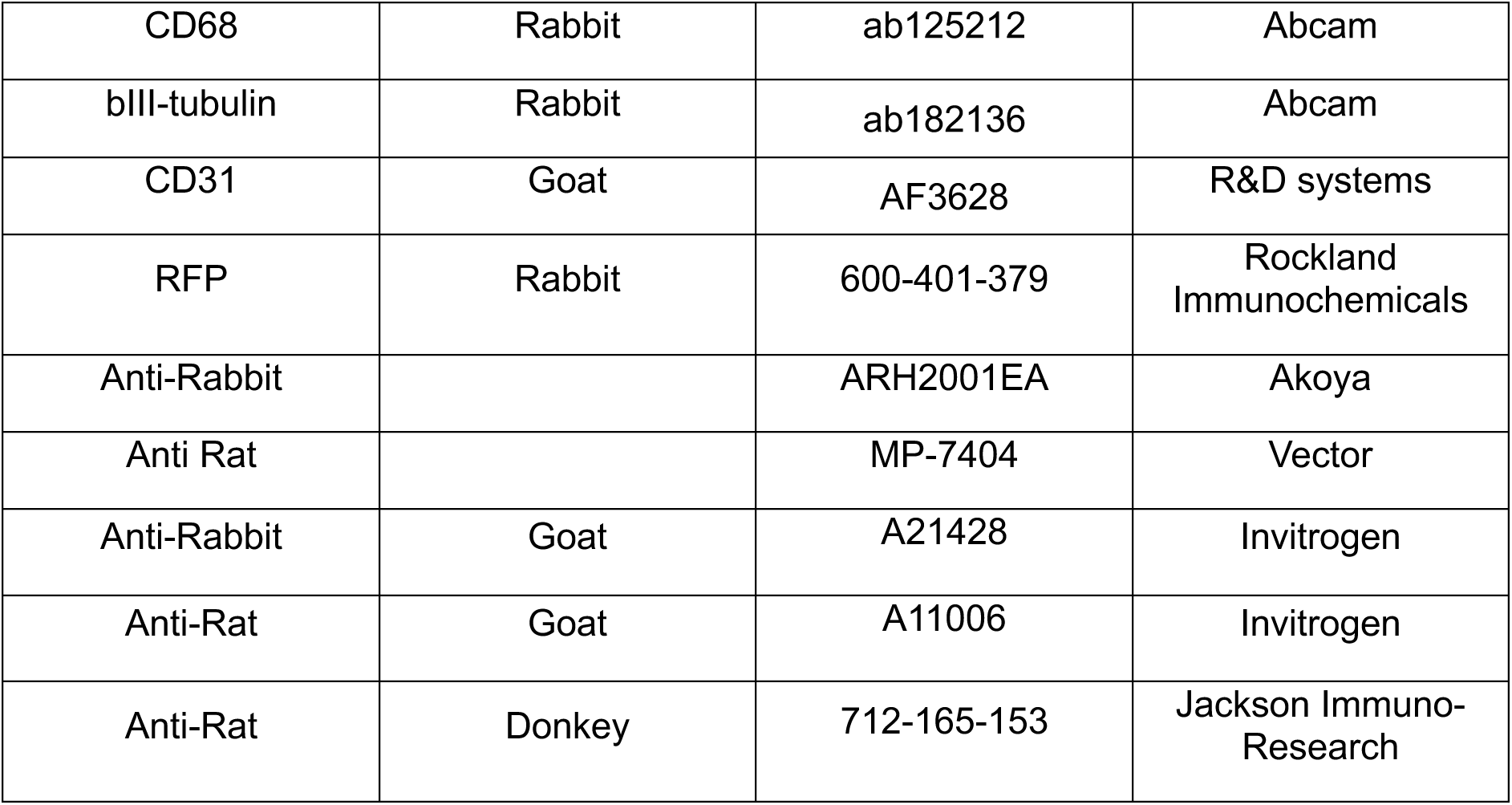
Antibodies used for immunofluorescence staining.

| **Antibody** | **Host species** | **Catalogue No.** | **Supplier** |
| --- | --- | --- | --- |
| CD11c | Hamster | 117312 | Biolegend |
| Ly6G | Rat | 551461 | BD Bioscience |
| CD68 | Rat | 137020 | Biolegend |
| EPCAM | Rat | 118222 | Biolegend |
| CD163 | Rat | 14-1631-82 | eBioscience |
| CD68 | Rabbit | ab125212 | Abcam |
| βIII-tubulin | Rabbit | ab182136 | Abcam |
| CD31 | Goat | AF3628 | R&D systems |
| RFP | Rabbit | 600-401-379 | Rockland<br>Immunochemicals |
| Anti-Rabbit |  | ARH2001EA | Akoya |
| Anti Rat |  | MP-7404 | Vector |
| Anti-Rabbit | Goat | A21428 | Invitrogen |
| Anti-Rat | Goat | A11006 | Invitrogen |
| Anti-Rat | Donkey | 712-165-153 | Jackson Immuno-Research |

### Imaging and quantification of immunofluorescence-stained tissue

Sections were imaged on a Zeiss LSM 780 confocal or a Zeiss Axioscan slidescanner. Cell numbers were quantified using the QuPath cell detection and cell classification tools[55]. 3×2 mm^2^ regions of interest (ROI) were chosen based on DAPI only. The ‘positive cell detection’ command was used to detect nuclei. Positive staining was then detected separately for each channel by using the ‘create single measurement classifier’ function. These classifiers were then combined with the ‘create combined classifier’ function. This allowed quantification of cells that had no positive antibody staining, ‘DAPI only’, and any combination of the other antibodies.

### Statistical analysis

Statistics were performed using Prism 10 (GraphPad Software). The statistical test used in each experiment is detailed in the relevant figure legend

## Supporting information

Combined Supplementary Data

## Acknowledgements.

Flow cytometry data were generated with the support from the IRR Flow Cytometry and Cell Sorting Facility, University of Edinburgh. Microscopy was supported by Shared University Research Facility (SURF) and the IRR Imaging Facility. We would like to thank Mariana Beltran and Elena Sutherland for technical expertise in setting up 10X sequencing and to Dr John Wilson-Kanamori for initial processing of the scRNA-seq raw data. Finally, we would like to thank the Bioresearch and Veterinary Services team at the University of Edinburgh for husbandry of our mice and other technical assistance.

## Author Contributions (CRediT)

L.H. conceptualisation, methodology, formal analysis, investigation, resources, data curation, writing - original draft/editing, visualisation, funding acquisition; GR.J. resources, writing - review/editing, funding acquisition; C.E.A. investigation, writing - review/editing. R.M.G resources. G.T.H. resources. E.E. resources, funding acquisition, writing - review/editing, supervision. C.C.B. conceptualisation, methodology, investigation, resources, data curation, writing - original draft/editing, visualisation, funding acquisition, supervision, project administration.

## Data and material availability

All data needed to evaluate conclusions in the paper are present in the paper or Supplementary Materials. Mouse scRNA-seq data will be deposited in the National Center for Biotechnology Information Gene Expression Omnibus public database (www.ncbi.nlm.nih.gov.geo/) upon acceptance for publication.

