## Supplementary material for "Tissue resident colonic macrophages persist through acute inflammation and adapt to aid tissue repair": Combined Supplementary Data

Supplementary Figure 1
Supplementary Figure 2
Supplementary Figure 3
Supplementary Figure 4

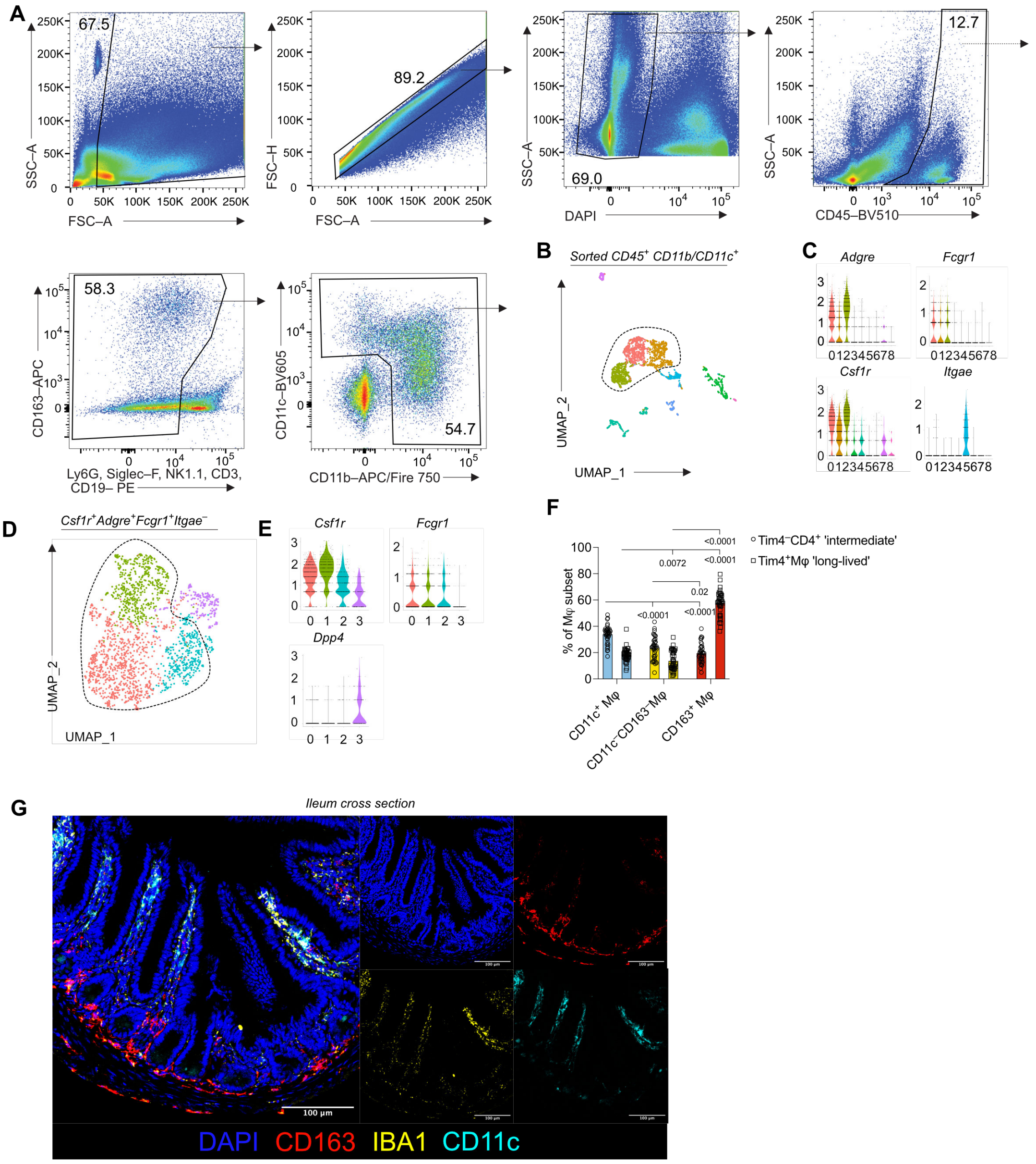

### Supplementary Figure 1

**A.** Gating strategy used for single cell RNA sequencing (scRNAseq): Forward scatter area (FSC−A) vs side scatter area (SSC−A) to exclude debris; FSC−A vs FSC−height (FSC-H) to exclude doublets/aggregates; SSC−A vs DAPI, live−dead marker to remove dead cells; SSC−A vs CD45 to select CD45+ leukocytes; CD163 vs Ly6G, SiglecF, NK1.1, CD3 and CD19 to exclude neutrophils, eosinophils, NK cells, T cells, and B cells respectively; CD11c vs CD11b to select mononuclear phagocytes which express CD11c or CD11b.

**B.** UMAP projection of sorted CD45$^{+}$ CD11c/CD11b$^{+}$ colonic cells from naïve adult C57BL/6 mice (as subclustered in Supplementary Fig. 1) and  Cells from three mice were pooled for analysis.

**C.** Violin plots of selected cluster-defining genes from **B.**

**D.** UMAP projection of *Csf1r/Fcgr1/Adgre1*$^{+}$*Itgae*$^{-}$ subclustered from **B.**

**E.** Violin plots of selected cluster-defining genes from **D.**

**F.** Mean frequency of macrophage subset that is Tim4$^{-}$CD4$^{+}$ (intermediate longevity), and Tim4$^{+}$CD4$^{+}$ 'long-lived'. Two-way ANOVA with Tukey's post hoc test. n=38 from across 7 individual experimentsin steady-state, C57bl/6J. Symbols represent individual mice. Both male and female mice are included and analysed together.

**G.** Representative immunofluorescence staining of Hoechst (blue), CD11c (cyan), IBA1 (yellow), CD163 (red) 20μm thick ileum Swiss roll sections.

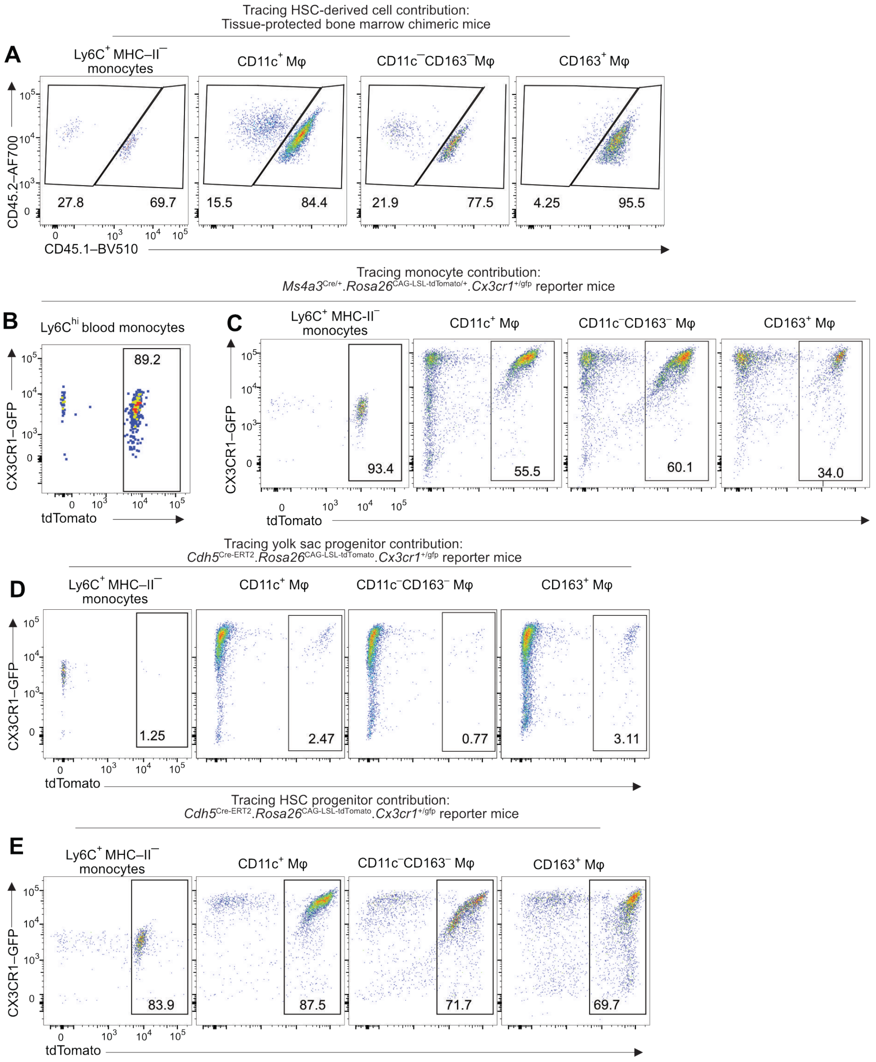


### Supplementary Figure 2

**A.** Representative flow cytometry plots of non-host Ly6C+ MHCII− monocytes, and CD11c+, CD163+, and CD11c−CD163− macrophages from tissue-protected BM chimeric mice, analysed 36 weeks after reconstitution.

**B.** Representative flow cytometry plot of CX3CR1−GFP and tdTomato expression by blood monocytes

(CD45$^+$CD11b$^+$CD115$^+$Ly6C$^{hi}$) in Ms4a3$^{Cre/+}$.Rosa26$^{LSL-CAG-tdTomato}$ lineage-tracing mice.

**C.** Representative flow cytometry plots of CX3CR1−GFP and tdTomato expression by Ly6C$^+$MHC-II$^-$ monocytes and CD11c$^+$, CD163$^+$, and CD11c$^-$CD163$^-$ macrophages in 12-week-old Ms4a3$^{Cre/+}$.Rosa26$^{LSL-CAG-tdTomato}$ mice.

**D.** Representative flow cytometry plots of CX3CR1−GFP and tdTomato expression by colonic Ly6C$^+$MHCII$^-$

monocytes and CD11c$^+$, CD163$^+$, and CD11c$^-$CD163$^-$ macrophages in *Cdh5*$^{Cre-ERT2/+}$.*Rosa26*$^{LSL-tdTomato/+}$ .*Cx3cr1*$^{+/GFP}$ fate mapping mice labelled at E7.5 analysed at 7 weeks of age.

**E.** Representative flow cytometry plots of CX3CR1−GFP and tdTomato expression by colonic Ly6C$^+$MHCII$^-$

monocytes and CD11c$^+$, CD163$^+$, and CD11c$^-$CD163$^-$ macrophages in *Cdh5*$^{Cre-ERT2/+}$.*Rosa26*$^{LSL-tdTomato/+}$ .*Cx3cr1*$^{+/GFP}$ fate mapping mice labelled at E10.5 analysed at 7 weeks of age.

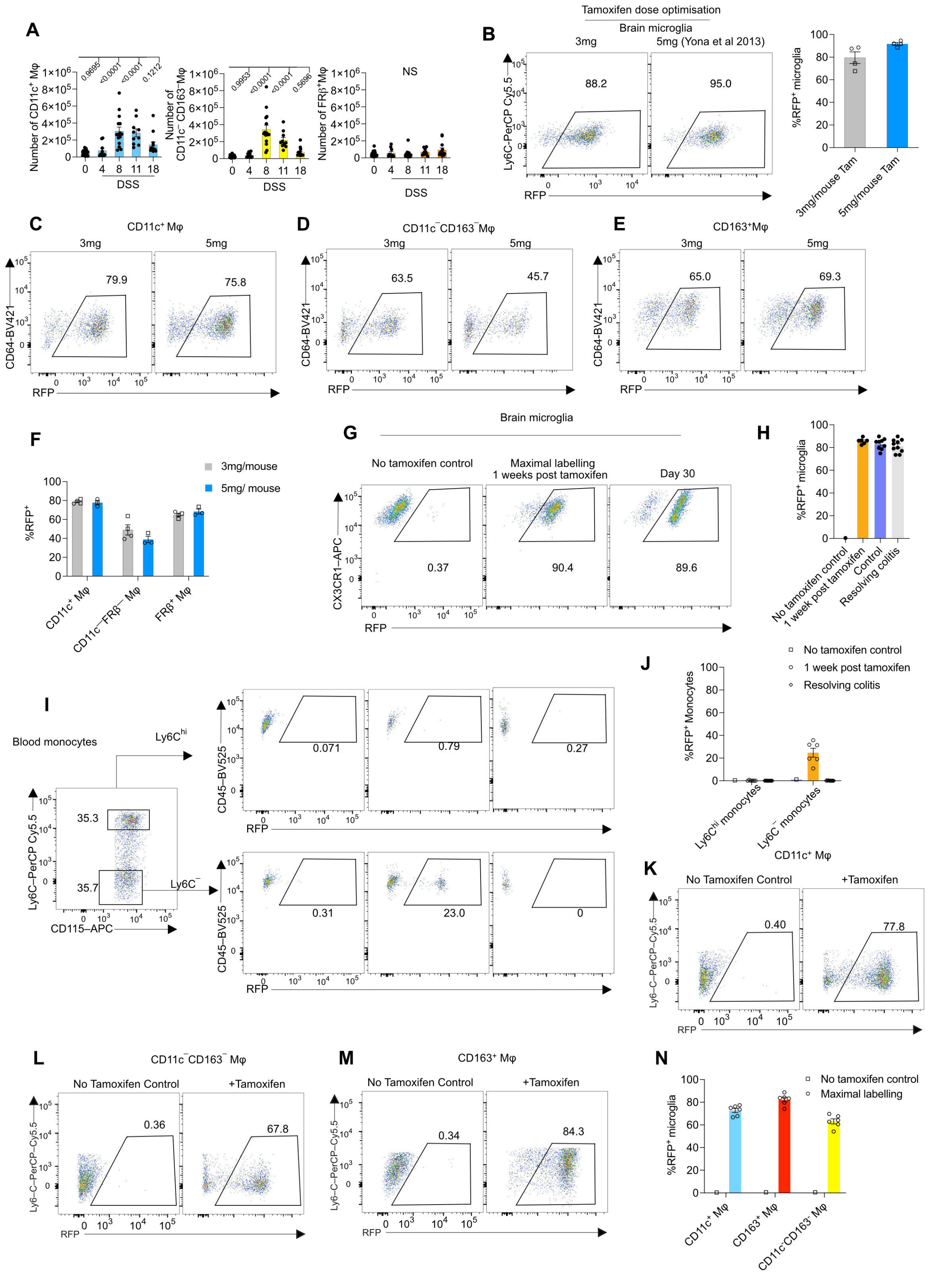


### Supplementary Figure 3

**A.** Mean number of CD11c⁺, CD11c⁻ FRβ⁻, and FRβ⁺ macrophages across the DSS time course. Data from three independent experiments with n=21 (control mice, day 0), n=15 (day 8), n=14 (day 18). All mice were male. One-way ANOVA followed by Dunnett's multiple comparisons test. Cell numbers are based on FlowJo cell counts, normalised to the numbers of Absolute Counting Beads™ detected per sample. Error bars ± SEM.

**B.** Representative flow cytometry plot, and mean expression of RFP by brain microglia following 3 or 5 mg tamoxifen administration every day for five days in *Cx3cr1*$^{Cre\text{-}ERT2/+}$.*Rosa26*$^{LSL\text{-}RFP/+}$ mice.

**C.** Representative flow cytometry plots of RFP expression by CD11c$^+$ macrophages following 3 or 5mg tamoxifen administration.

**D.** Representative flow cytometry plots of RFP expression by CD11c$^-$ CD163$^-$ macrophages following 3 or 5mg tamoxifen administration.

**E.** Representative flow cytometry plots of RFP expression by CD163$^+$ macrophages following 3 or 5mg tamoxifen administration.

**F.** Mean RFP expression following 3 or 5mg tamoxifen administration.

**G.** Representative flow cytometry plots of RFP expression by brain microglia (CD45$^{int}$CD11b$^+$MHCII$^-$CX3CR1$^+$) obtained from *Cx3cr1*$^{Cre\text{-}ERT2/+}$.*Rosa26*$^{LSL\text{-}RFP/+}$ mice one week (maximal labelling) or 29 days post tamoxifen administration, or no tamoxifen controls.

**H.** Mean RFP expression by brain microglia (CD45$^{int}$CD11b$^+$MHCII$^-$CX3CR1$^+$).

**I.** Representative flow cytometry plots of RFP expression by blood monocytes (CD45$^+$CD11b$^+$ CD115$^+$) obtained from *Cx3cr1*$^{Cre\text{-}ERT2/+}$.*Rosa26*$^{LSL\text{-}RFP/+}$ mice one week (maximal labelling) or 29 days post tamoxifen administration, or no tamoxifen controls.

**J.** Mean expression of RFP by blood monocytes (CD45$^+$CD11b$^+$ CD115$^+$). Data are pooled from two independent experiments with n=1 (No tamoxifen control), n=6 (Maximal labelling), n=9 (Control) and n=10 (resolving colitis). Both male and female mice were used and analysed together. Symbols represent individual mice. No statistical analysis was performed. Error bars ± SEM.

**K.** Representative flow cytometry plots of RFP expression by CD11c$^+$ macrophages one-week post tamoxifen.

**L.** Representative flow cytometry plots of RFP expression by CD11c$^-$ CD163$^-$ macrophages one-week post tamoxifen.

**M.** Representative flow cytometry plots of RFP expression by CD163$^+$ macrophages one-week post tamoxifen.

**N.** Mean RFP expression. Symbols represent individual mice. Data are pooled from two independent experiments with n=1 (no tamoxifen control), n=6 (maximal labelling). Both male and female mice were used in this experiment. No statistical analysis was performed. Error bars ± SEM.

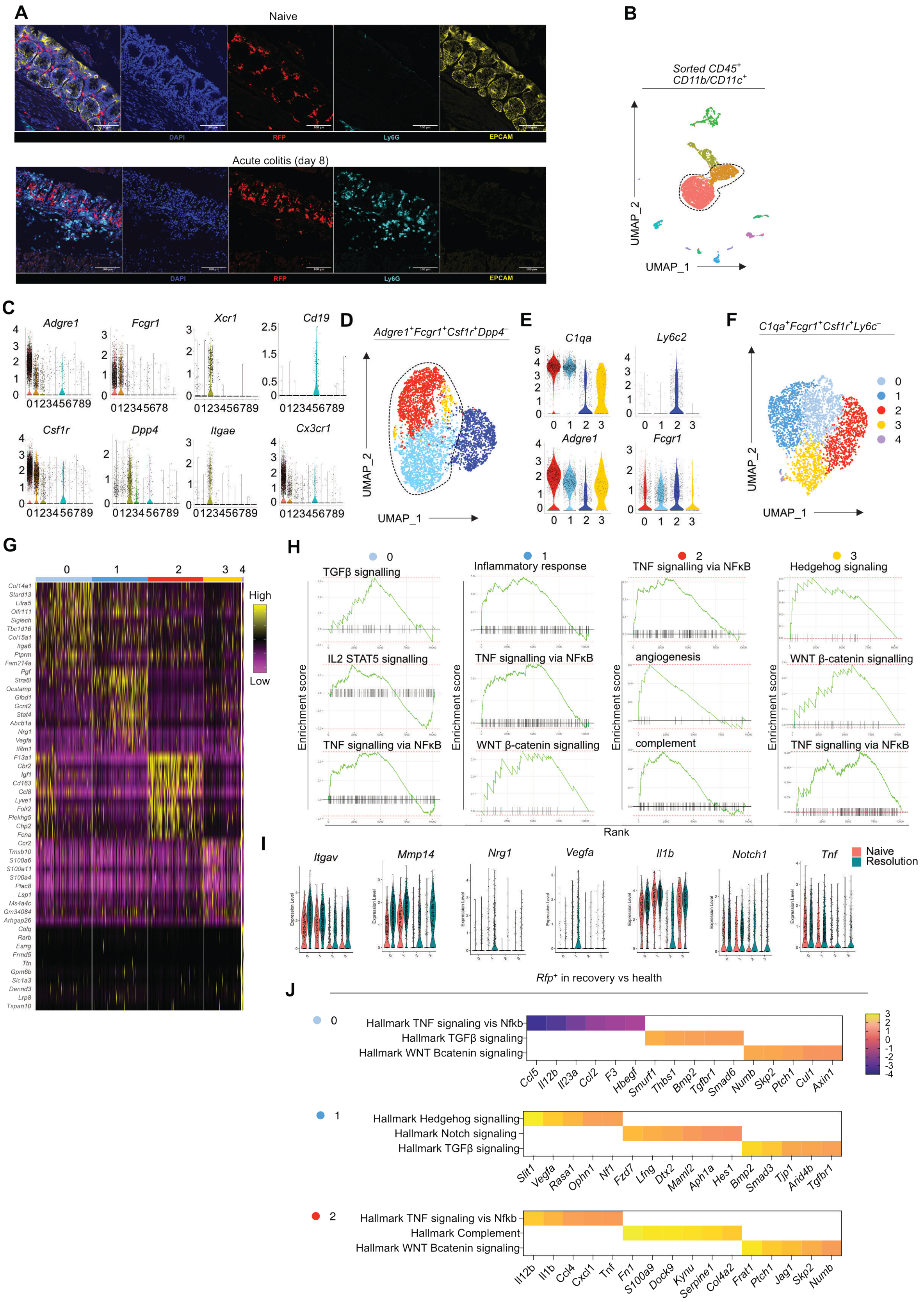


### Supplementary Figure 4

**a.** Immunofluorescence staining of DAPI (blue), RFP (Red; representing fate-mapped macrophages), Ly6G (cyan; marking neutrophils) and EPCAM (yellow; marking epithelial cells) in tamoxifen treated naïve (A) or acutely colitic (B) *Cx3cr1*$^{\text{Cre-ERT2/+}}$.*Rosa26*$^{\text{LSL-RFP/+}}$ mice. 12μm thick Swiss roll sections. Data are from day 8 of the DSS time-course (following 4 days 2% DSS treatment, and 4 days normal drinking water). Images are representative of n=4 naïve n=5 acutely colitic male and female mice.

**B.** UMAP projection of sorted CD45$^{+}$ CD11c/CD11b$^{+}$ colonic cells from naïve and resolution *Cx3cr1*$^{Cre\text{-}ERT2/+}$.*Rosa26*$^{LSL\text{-}RFP/+}$ mice mice.

**C.** Violin plots of selected cluster-defining genes from **B.**

**D.** UMAP projection of *Csf1r/Fcgr1/Adgre1*$^{+}$*Itgae/Cd19/Xcr1/Dpp4*$^{-}$ cells subclustered from **B.**

**E.** Violin plots of selected cluster-defining genes from **D.**

**F.** UMAP (*C1qa*$^{+}$*Ly6c2*$^{-}$) macrophages sub-clustered from **D**.

**G.** Heatmap displaying the top 10 cluster defining genes for each cluster of macrophages in **F.**

**H.** Fast gene set enrichment analysis (FGSEA) using Hallmark gene set signatures of each cluster in resolution vs naïve. Enrichment plots.

**I.** Violin plots of selected genes in resolution vs. naive.

**J.** Fast gene set enrichment analysis (FGSEA) using Hallmark gene set signatures of the *Rfp*$^{+}$ macrophages in each cluster in resolution vs naïve. Heatmap of leading-edge genes for each pathway.
